# Optogenetic Targeting of AII Amacrine Cells restores Retinal Computations performed by the Inner Retina

**DOI:** 10.1101/2022.07.28.501925

**Authors:** Hanen Khabou, Elaine Orendorff, Francesco Trapani, Marco Rucli, Melissa Desrosiers, Pierre Yger, Deniz Dalkara, Olivier Marre

## Abstract

Most inherited retinal dystrophies display progressive photoreceptor cell degeneration leading to severe visual impairment. Optogenetic reactivation of inner retinal neurons is a promising avenue to restore vision in retinas having lost their photoreceptors. Expression of optogenetic proteins in surviving ganglion cells, the retinal output, allows them to take on the lost photoreceptive function. Nonetheless, this creates an exclusively ON retina by expression of depolarizing optogenetic proteins in all classes of ganglion cells, whereas a normal retina extracts several features from the visual scene, with different ganglion cells detecting light increase (ON) and light decrease (OFF). Refinement of this therapeutic strategy should thus aim at restoring these computations. In an attempt to do so, we used a promoter that targets gene expression to a specific interneuron of the retina called the AII amacrine cell. The AII amacrine cell simultaneously activates the ON pathway and inhibits the OFF pathway. We show that the optogenetic stimulation of AII amacrine cells allows restoration of both ON and OFF responses in the retina, but also mediates other types of retinal processing such as sustained and transient responses. Targeting amacrine cells with optogenetics is thus a promising avenue to restore better retinal function and visual perception in patients suffering from retinal degeneration.

## Introduction

Blindness affects 45 millions people worldwide. In many cases of retinal degeneration photoreceptors are lost, while retinal ganglion cells (that provide visual signals to the brain) as well as many interneurons (e.g. amacrine cells) are spared. This opens the possibility to stimulate the remaining retinal ganglion or amacrine cells directly to restore visual function. Retinal prostheses are a promising solution and have been found to restore some useful perception in blind patients. However, the acuity of the existing devices remains very low, below the level of legal blindness [Lorach et al. 2012, da Cruz et al. 2016]. Patients also report that percepts evoked by electrical stimulation of retinal neurons are not easily interpretable as visual stimuli [Beyleler et al. 2019] and therefore are often not sufficient to identify objects or to navigate in complex environments. Optogenetic therapies provide a possible alternative to restore vision with a higher resolution and specificity that can better mimic the natural output of the retina [Ferrari et al. 2020]. In this strategy, a light sensitive protein is expressed in specific neural populations in a blind retina.

Expressing light sensitive proteins in retinal ganglion cells can be an efficient way to restore vision through the stimulation of these newly light-sensitive cells with patterned light to evoke visual perception [Bi et al. 2006, Caporale et al. 2011, Sengupta et al. 2016, Chaffiol et al. 2017, Berry et al. 2019], although the first results show that the acuity is still low with this strategy [Sahel et al. 2021]. How to optimize visual acuity and perceptual performance when restoring vision using optogenetics is an active area of investigation.

In a healthy retina, the ganglion cell population can be divided into about 20 to 40 cell types that each perform a different computation on the visual scene [Sanes & Masland 2015, Baden et al. 2016]. Each cell type is classically assumed to be selective of a specific feature of the visual scene and therefore convey a corresponding feature map to the brain [Deny et al. 2017]. Altering specifically one of these cell type populations can lead to specific impairments in visual perception and motor output, including specific defects in perceiving moving objects and eye movement control [Merigan et al. 1991, Yonehara et al. 2016, Hillier et al. 2017]. In particular ganglion cells usually respond either to light increase (ON ganglion cells) or light decrease (OFF ganglion cells). Inactivating ON ganglion cells leads to a reduced ability to detect increase of luminance at the perceptual level while ability to detect decrease of luminance is not affected [Schiller et al. 1986]. Optogenetic strategies targeting ganglion cells will not restore the computations performed in the normal retina. In particular, making ganglion cells light-sensitive will result in a retina where all ganglion cells become de facto ON cells (only responding to light increase). It is unclear how this synthetic visual signal will affect the physiological processing performed by downstream areas in the brain and what will be the resulting restored perception, but this loss of retinal computations could severely impair perceptual performance.

To restore some of the response diversity found in normal retinas with optogenetic therapy, other cells such as “dormant” photoreceptors have been targeted specifically [Busskamp et al. 2010]. However in many patients these cells are not viable targets, as they are also affected by retinal degeneration. An alternative strategy to restore richer functional selectivity is to target other cell types in the intermediate layers of the retina that are not damaged. Here we express an optogenetic protein in AII amacrine cells of blind mice to restore vision. AII amacrine cells are an ideal target because they connect with both ON and OFF bipolar cells with different types of synapses. They form gap junctions with most ON bipolar cell types [Tsukamoto & Omi 2017], and can therefore excite them when they are activated. At the same time, they form glycinergic inhibitory synapses with most OFF bipolar cell types [Tsukamoto & Omi 2017].

We first introduce a promoter allowing to target specifically AII amacrine cells, enabling expression of an optogenetic protein following an AAV injection. We then show that this strategy allows reactivating retinal computations, and in particular ON-OFF selectivity, in a way similar to the normal retina. We demonstrate this both in normal retinas where photoreceptor transmission is blocked and in a model of retinal degeneration where photoreceptors have been lost. Our data shows that targeting AII amacrine cells is a promising strategy for vision restoration with optogenetics.

## Results

### Targeting of AII Amacrine Cells

To identify a promoter that can drive expression in AII amacrine cells, we tested several known retinal specific promoters via intravitreal injection route encapsidated in AAV2-7m8, a genetic variant of AAV2 [Dalkara et al. 2013, Khabou et al. 2018]. We incidentally found a sequence driving specific expression in AII amacrine cells, that we refer to as HKamac in the following (see supplementary material for the sequence). 5 mouse eyes of C57BL/6J wild-type were injected intraocularly at 4 weeks of age using this promoter driving GFP expression in AAV2-7m8 capsid. 6 weeks after injection, eye fundus showed high expression levels. Retinas were then harvested, fixed, and embedded in tissue freezing medium for histology and confocal microscopy 120–140 days postinjection.

Wild-type retinal flat mounts showed that a homogeneous population of cells with large somas expressed GFP in the INL (fig. 1.A). In cross-sections, the labeled cells showed dendritic stratification in both ON and OFF layers, a pattern reminiscent of AII morphology [Helmstaedter et al. 2013]. To determine precisely the subtype of amacrine cell, we first showed that GFP did not colocalize with GABAergic or with Starbust amacrine cell marker (fig. 1.B) but did colocalize with a glycinergic cell marker. To further confirm that they were AII amacrine cells, we performed cryosections labelings with Prox1, an antibody labeling both glycinergic bipolar cells and AII amacrine cells, and found clear co-localization of GFP with Prox1, thereby confirming GFP positive cells are AII amacrine cells (Fig. 1.C, D, E). We then replaced GFP by ReaChR in the plasmid construct while keeping the same promoter. ReachR was then delivered in wild-type mice using the same AAV2-7m8 capsid variant. ReachR was successfully expressed in AII amacrine cells, although it was detected in some RGCs as well (fig. 1.F, G, H).

**Figure 1:**
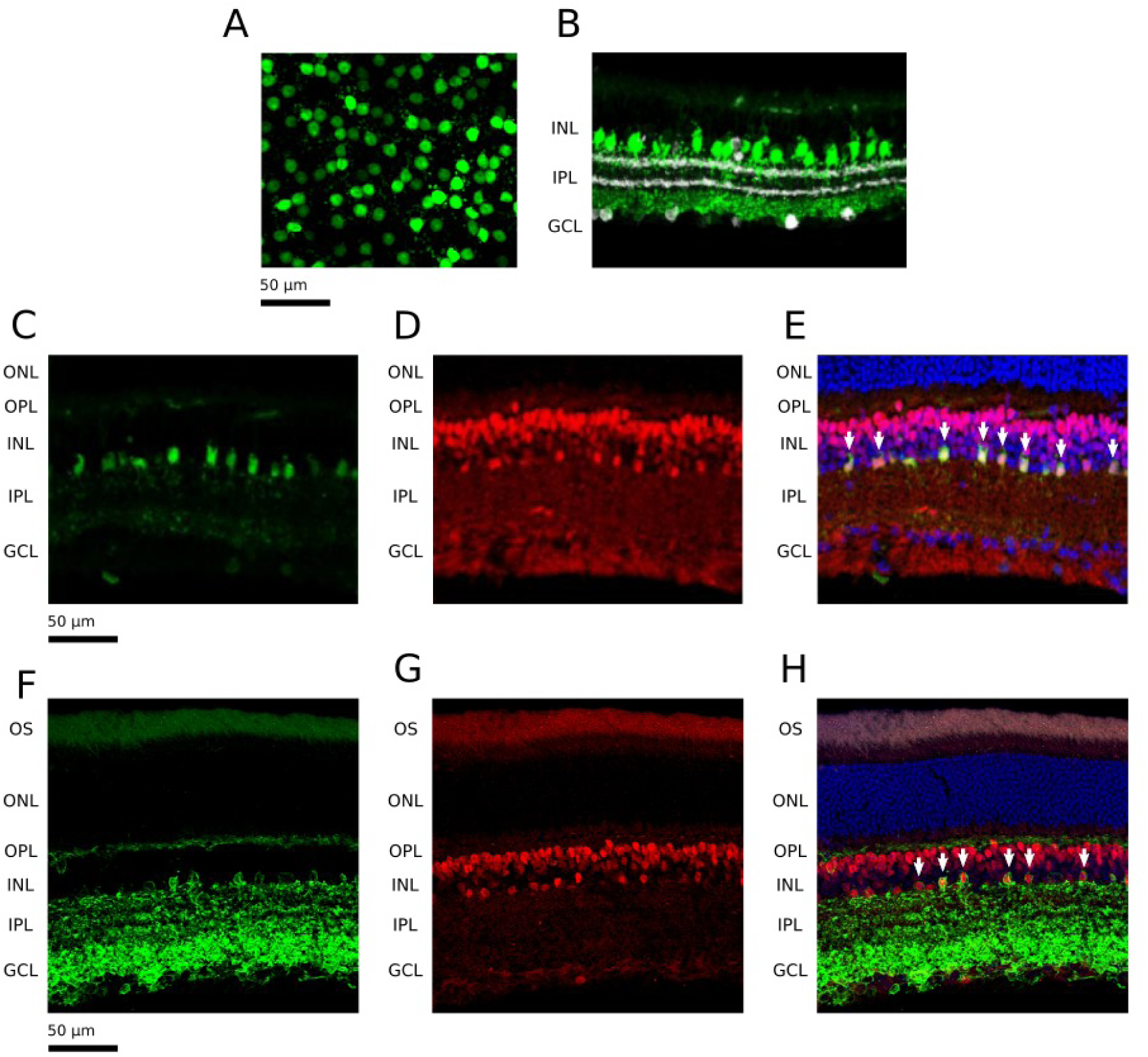
AAV-mediated gene delivery to AII amacrine cells from the vitreous. **A**. Whole-mount view showing GFP expression (green) in the plane of AII somas. **B**. Side-view showing GFP expression (green) and labeling of starburst amacrine cells with a ChAT antibody (white). **C**,**D**,**E**. Side-view showing GFP expression respectively under the control of: our promoter (in green, panels C and E), Prox1 antibody (in red, panels D and E) and DAPI (in blue, panel E). DAPI labels all nuclei, while Prox1 labels bipolar and AII amacrine cells. AII amacrine cells are indicated with white arrows. **F**,**G**,**H**. Side-view showing ReachR expression respectively under the control of: our promoter (in green, panels F and H), Prox1 antibody (in red, panels G and H) and DAPI (in blue, panel H). AII amacrine cells are indicated with white arrows. For all panels, retinal layers are shown on the left: outer segment (OS), outer nuclear layer (ONL), outer plexiform layer (OPL), inner nuclear layer (INL), inner plexiform layer (IPL), ganglion cell layer (GCL).

### Optogenetic Stimulation Of AIIs Produces On And Off Responses

AII amacrine cells excite ON bipolar cells through gap junction, and inhibit OFF bipolar cells through glycinergic, inhibitory synapses. We tested if stimulation of AII amacrine cells with optogenetics could evoke ON and OFF responses in retinal ganglion cells (RGCs). For this we recorded RGC spiking activity from wild-type retinas expressing ReachR under the HKamac promoter on a multielectrode array (fig. 2.A). To test if AII stimulation could activate similar circuits to photoreceptor stimulation, we first measured the responses of ganglion cells to stimuli at low light intensity, which only activated photoreceptors, and were not sufficiently strong to activate ReachR (termed photoreceptor stimulation in the following). We observed both ON and OFF responses (fig. 2.B). We then blocked the synaptic transmission from photoreceptors to the ON and OFF bipolar cells using pharmacology (L-AP4 to block the transmission from photoreceptors to ON bipolar cells, and ACET to block transmission from photoreceptors to OFF bipolar cells, see methods). At the same light intensity, responses disappeared. This is expected since the impact of photoreceptor activation on the rest of the retinal circuit has been blocked, and the light intensity is too low to activate ReachR.

**Figure 2:**
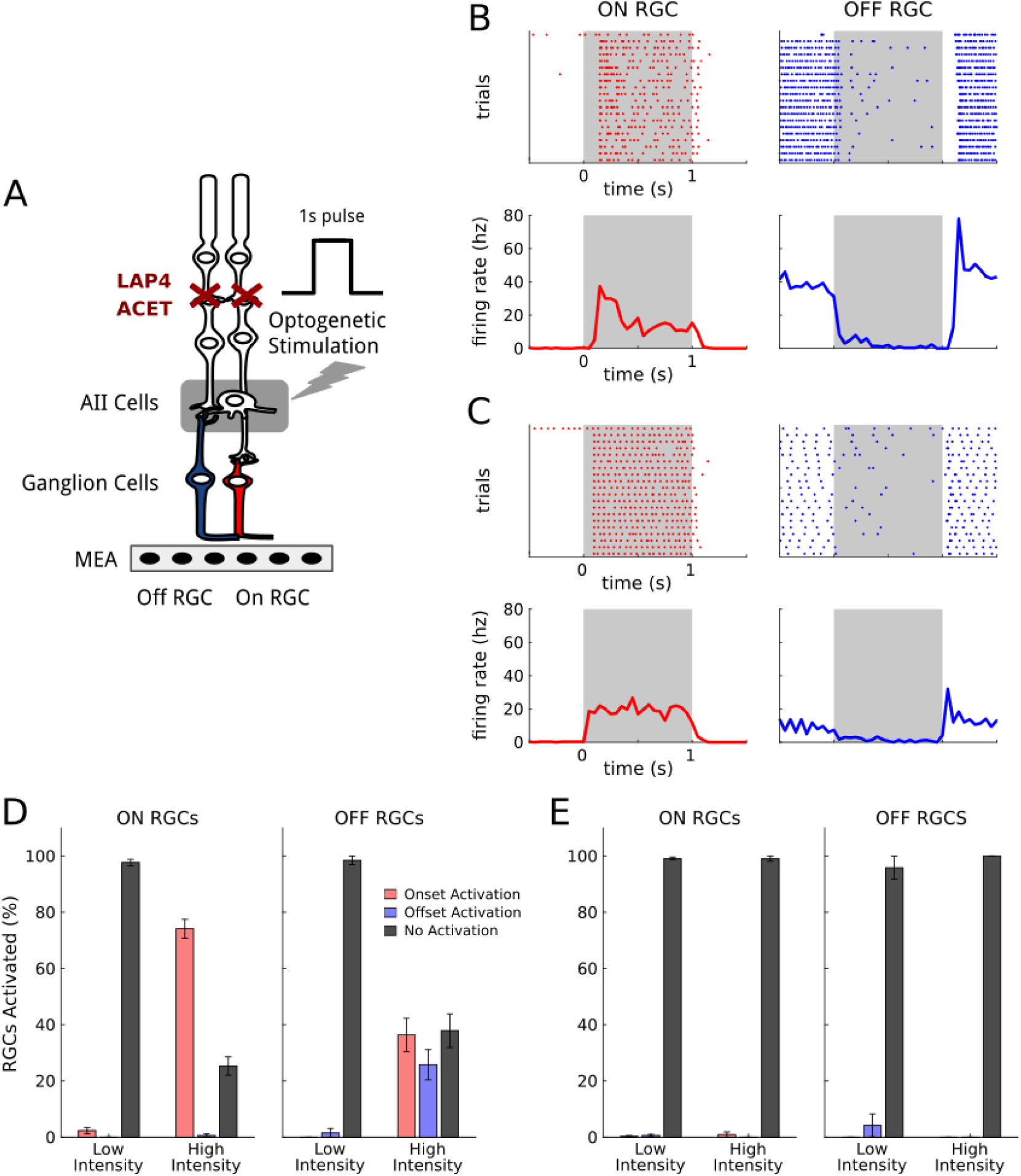
ON and OFF RGC responses elicited upon light stimulation of ReaChR-expressing AII amacrine cells. A. AII amacrine cells connect to the ON pathway (in red) through gap junctions, and to the OFF pathway (in blue) through glycinergic inhibitory connections. Stimulation of AIIs hence produces responses of opposite polarity on ON and OFF retinal ganglion cells. We target AIIs through optogenetic stimulation consisting of a series of full field flashes, and record the responses of the retinal ganglion cells with a multielectrode array. Pharmacology (ACET and L-AP4) blocks synaptic transmission from photoreceptors. B. Responses of representative ON (left column, red) and OFF (right column, blue) retinal ganglion cells to photoreceptor stimulation with white full-field flashes. Top: spiking activity across different trials. Bottom: mean response. The time intervals of the flashes are shown in gray. C. Responses of the same ON (left column, red) and OFF (right column, blue) RGCs shown in B to optogenetic stimulation, after blocking photoreceptor synaptic transmission. Top: spiking activity across different trials. Bottom: mean response. D. Percentage of retinal ganglion cells responding to optogenetic stimulation. Left: percentage of ON RGCs responding to the flash respectively at onset (red), offset (blue), or never (black), for both low and high luminance. Right: same plot for OFF RGCs. E. Percentage of retinal ganglion cells responding to optogenetic stimulation for a control population with no opsin expressed. Same plots as in D.

We then increased light intensity to reach the activation of ReachR (see methods) and observed both ON and OFF responses to light stimulation (fig. 2.C, D) for a large fraction of ganglion cells. This activation (termed optogenetic stimulation in the following) is due to the stimulation of AII since stimulation at similar intensity in control retinas with the same concentration blockers, but no AAV injection, did not show any response (fig. 2.E). Activation of AII with optogenetic stimulation is thus able to evoke both ON and OFF responses in the retina.

To understand if our strategy allows reactivating the same computations performed in the normal retina, we categorized the cells as ON or OFF depending on their responses to photoreceptor stimulation. ON ganglion cells were defined as responding to the onset of the photoreceptor stimulation, and OFF cells as responding to the offset (see methods).

We then asked if ON and OFF cells also responded to the onset and offset of the optogenetic stimulation respectively. If the responses were consistent for photoreceptor and optogenetic stimulation, this would suggest that AII stimulation is able to reactivate some of the circuits that are active during photoreceptor stimulation. For example, ON ganglion cells receive their inputs from ON bipolar cells. At the onset of AII stimulation, ON bipolar cells should be activated, and should therefore stimulate ON ganglion cells. Conversely, OFF bipolar cells, which provide the main excitatory input to OFF ganglion cells, should be inhibited during AII stimulation, and disinhibited at the offset of AII stimulation. As a result, they should be able to excite OFF ganglion cells at the offset of the AII stimulation. If this hypothesis is correct, ON ganglion cells should be activated at the onset of AII stimulation, and OFF ganglion cells at the offset.

We identified a total population of 173 ON and 65 OFF ganglion cells across 3 different experiments. The large majority of ON cells responded to the onset of the optogenetic stimulation (74%) while their responses to the offset were almost not present (only 0.5%), which is consistent with our hypothesis. A significant portion of OFF cells showed OFF responses to the optogenetic stimulation (25%). However, we also observed that a fraction of cells classified as OFF based on response to photoreceptor stimulation, turned ON for optogenetic stimulation (36%). We hypothesized that this could be due to off-target expression of the ReachR protein, observed in our histology experiments.

### Off-target Expression Explains Changes In ON-OFF Selectivity

If the observed responses at the onset for OFF cells are due to direct expression of ReachR in ganglion cells, these responses should still be present when fully blocking glutamatergic synaptic transmission. To test if this was the case, we performed additional tests on the same cell populations where we fully blocked this transmission using a pharmacological cocktail composed of L-AP4, ACET, CNQX and CPP (fig. 3.A, see methods). The responses at the stimulation onset in OFF ganglion cells were still present after the application of this cocktail (fig. 3.B, C, D): 41% of the OFF cell population responded to the stimulus onset at bright luminance, while responses to the stimulus offset completely disappeared (fig. 3.E). In control retinas with no optogenetic protein expressed, instead, light responses were almost completely abolished (99% or the OFF population didn’t respond to the stimulus, fig. 3.F). This confirms that these onset responses in OFF cells are due to off-target expression in ganglion cells.

**Figure 3:**
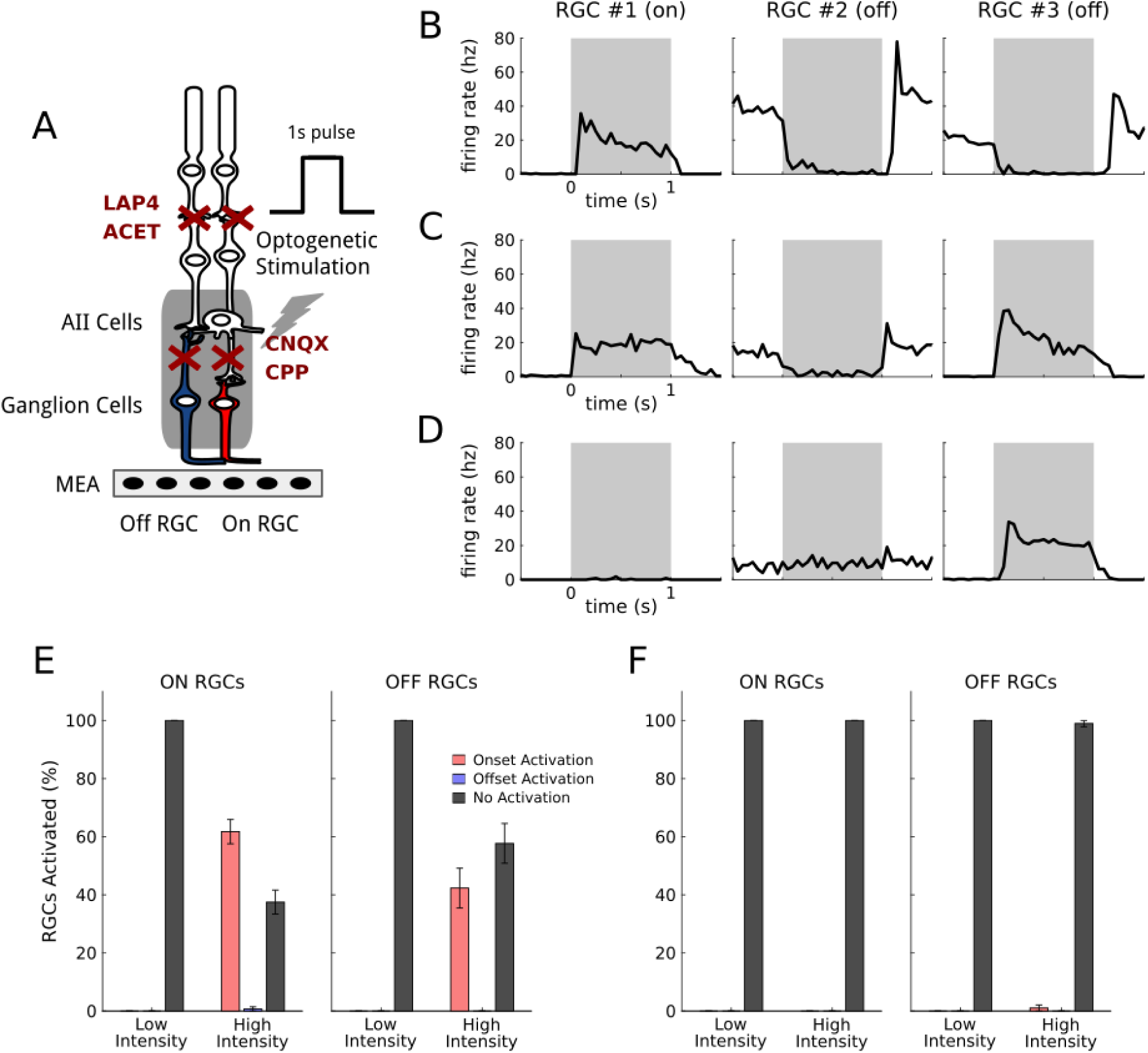
Off RGC Responses of inverted polarity are due to off-target opsin expression. **A**. Control protocol showing direct ganglion cell activation due to off-target expression of ReaChr: the application of CNQX and CPP disrupts all the excitatory synaptic connections. Responses induced by visual stimulations under this condition are only due to direct activation of the retinal ganglion cells expressing the opsin. **B**. Mean responses of three representative RGCs (one ON and two OFF) to simple photoreceptor stimulation. The stimulation time interval is depicted in gray. **C**. Mean responses of the same three RGCs to optogenetic stimulation, after blocking the photoreceptor transmission. **D**. Mean responses of the same RGCs after blocking all the excitatory synaptic connections. Responses of RGC 3 can only be explained by off-target expression of the opsin in ganglion cells. **E**. Percentage of ganglion cells responding to direct optogenetic stimulation, due to off-target expression. Left: percentage of ON RGCs responding respectively to the stimulus onset or offset (or not responding), at both low and high luminance. Right: same plot for OFF RGCs. **F**. Percentage of ganglion cells responding to direct optogenetic stimulation for a control population with no opsin expressed. Same plots as in E.

### Ganglion Cell ON And OFF Responses Can Be Evoked By AII Stimulation

To further demonstrate that the observed responses were mostly due to AII activation and not off-target expression in ganglion cells, nor to a failure of the pharmacological blocking of photoreceptor transmission, we performed additional experiments. We reasoned that if we use an inhibitory opsin, which will hyperpolarize the cells upon light stimulation, it will inactivate ganglion cells. As a consequence, off-target expression will not allow any spiking response. On the contrary, if AII are hyperpolarized upon light stimulation, they should still evoke responses, except that they should be inverted: ON ganglion cells should respond at light offset, and OFF ganglion cells at light onset. If the responses are due to a failure of the pharmacological blocking, we should not see this inversion.

We injected the same construct but replaced ReachR with gtACR1 [Govorunova et al. 2015] (see methods). We performed the same protocols (2 retinas, 36 ON and 125 OFF RGCs) and found that most ganglion cells for which a response was detected showed the predicted inversion of polarity (fig. 4.A, B, C). Observed responses are thus due to AII modulation, and not to off-target expression, nor to a failure of the pharmacological block. These results confirm that our approach allows stimulating AII to modulate differentially ON and OFF ganglion cells.

**Figure 4:**
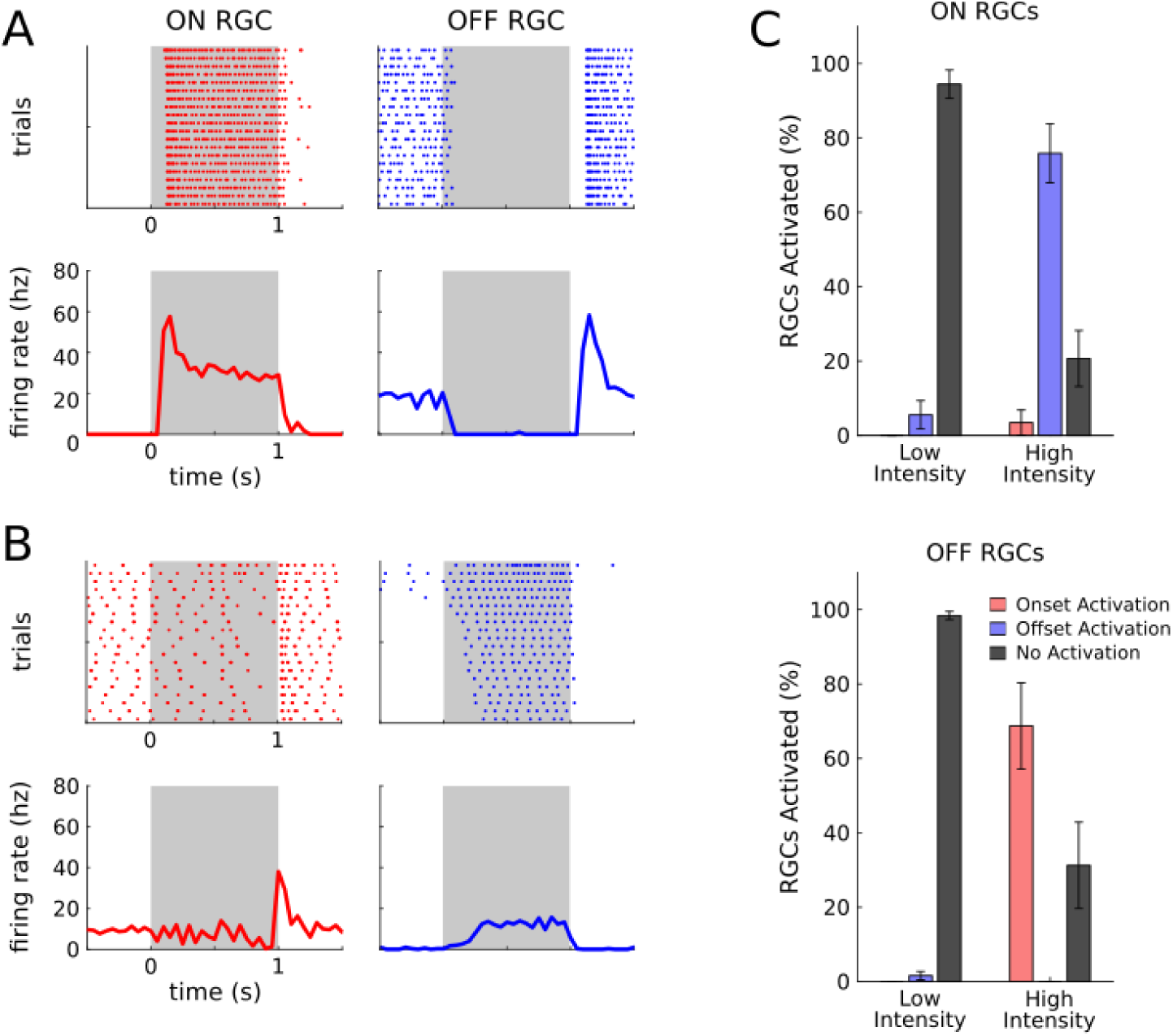
The diversity of RGC responses is really due to AII activation, and not to photoreceptor transmission or off-target expression. **A**. Examples of responses to photoreceptor stimulation for a representative ON (left column, red) and OFF (right column, blue) retinal ganglion cells. Top: raster plot of RGC responses across trials. Bottom: mean responses. The stimulation period is represented by the gray regions. **B**. same as A for optogenetic stimulation, after blocking photoreceptor transmission: this time AII amacrine cells express an inhibitory opsin, gtACR. **C**. Percentage of RGCs responding to optogenetic stimulation (with the inhibitory opsin gtACR). Top: percentage of ON RGCs responding respectively at stimulus onset (red), stimulus offset (blue), or not responding (black), for both low and high luminance. Bottom: same plot for OFF RGCs.

### Diversity Of Ganglion Cell Responses To AII Stimulation

AII stimulation can restore ON and OFF responses, but can it restore more features of the normal retinal responses? In particular, beyond the ON and OFF classification, previous works on normal retinas have shown that retinal responses to more complex stimuli like “chirp” stimuli uncover a large diversity of responses, corresponding to the different types of ganglion cells. Is this diversity still present in retinas reactivated by our AII stimulation strategy? To test this, we displayed the chirp stimulus, previously used to classify different types of ganglion cells [Baden et al. 2016], at high light intensity so that it activates the ReachR protein expressed in AII amacrine cells. We found a large diversity of responses to this stimulus (fig. 5.A). Beyond responses to onset and offset, some cells responded to different parts of this stimulus, showing different tunings to temporal frequencies.

**Figure 5:**
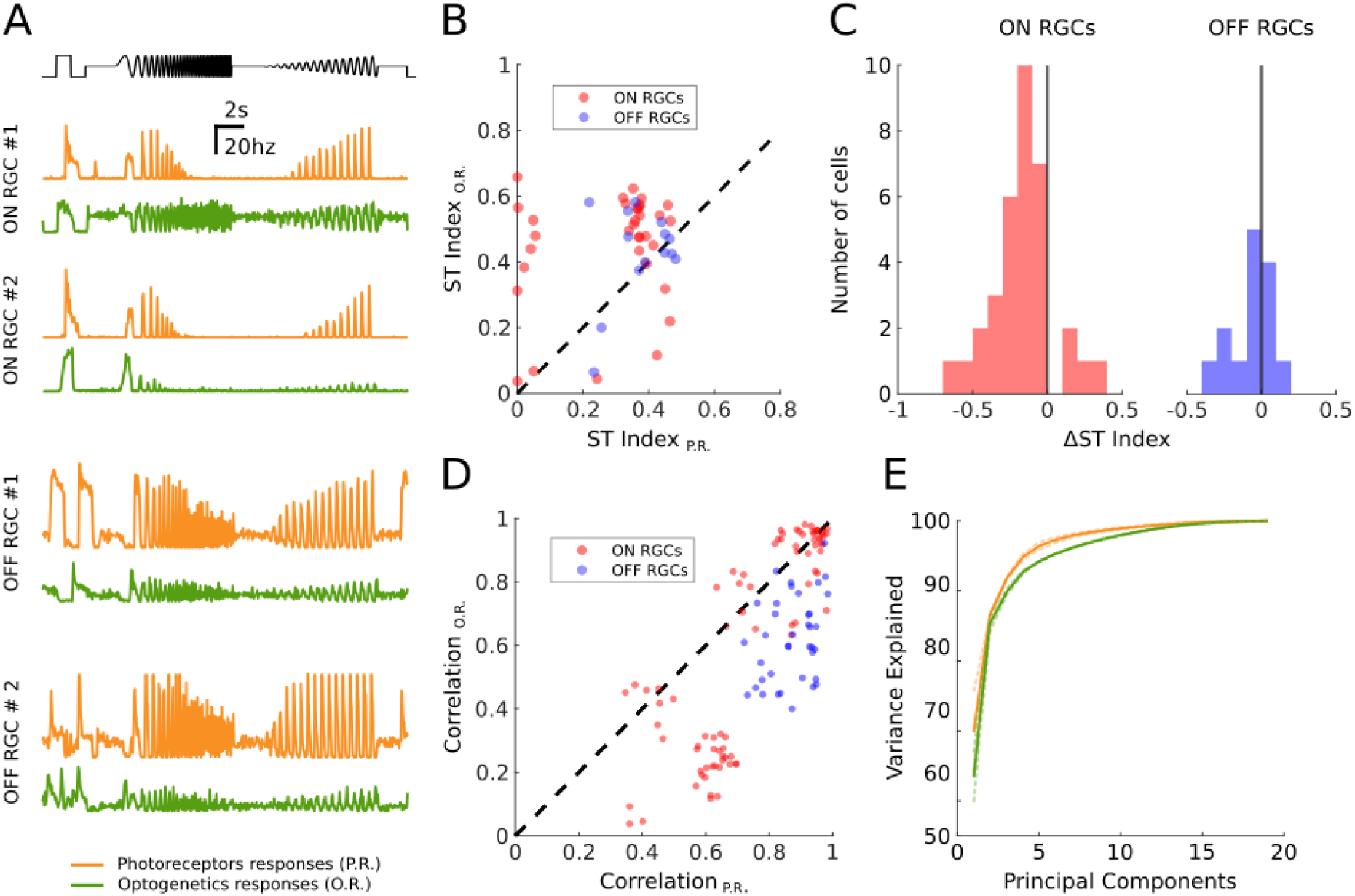
AII activation generates diverse RGC response. **A**. On top in black: the temporal profile of the full-filed chirp stimulus (luminance over time). Below: mean responses of four representative retinal ganglion cells to the chirp stimulus. In orange: responses mediated by photoreceptors (no optogenetics involved). In green: responses mediated by the optogenetic activation of AIIs (with blocked photoreceptor transmission) **B**. Comparison between the Sustained-Transient index computed on photoreceptor and optogenetic responses for 48 selected RGCs. Each dot shows the index for a different RGCC, for both photoreceptor responses (X axis) and optogenetic responses (Y axis). Values close to one mean the response is predominantly sustained; values close to zero mean the response is predominantly transient. **C**. Distribution of the comparative sustained-transient index ΔST for the ON (red) and OFF (blue) RGC populations (same 48 RGCs shown in B). An indexST for the ON (red) and OFF (blue) RGC populations (same 48 RGCs shown in B). An index close to 1 means that the cell responds more transiently when activated optogenetically with respect to its normal photoreceptor responses. Conversely, an index close to -1 means that the optogenetic responses are more sustained than the normal photoreceptor responses. **D**. Response correlations across retinal ganglion cell pairs from a population of 40 RGCs, for both photoreceptor and optogenetic stimulations. Pearson correlation coefficient computed on the responses of pairs of RGCs to photoreceptor stimulation (X axis) and to optogenetic stimulation (Y axis). Red dots represent ON to ON response correlations: blue dots represent OFF to OFF response correlations. **E**. Principal Component Analysis of the mean RGC responses to the chirp stimulus for a selected population of 40 cells (same shown in D). Each curve shows the number of principal components needed (X axis) to explain a given percentage of variance in the ganglion cell responses (Y axis). We show the curves for both photoreceptor responses (orange) and optogenetic responses (green). Light dashed curves represent analysis conducted on the individual experiments. Dark, continuous lines represent the average across all experiments.

A few cells responded transiently while most of them had more sustained responses. To quantify this, we first measured an index of how transient or sustained ganglion cells responses were (see methods). We computed this index both on the responses to optogenetic stimulation and to normal photoreceptor responses: we found that a large majority of cells lost their transient component when reactivated through the AII pathway (fig. 5.B). We also defined a comparative sustained-transient index, that measures how transient is the response of a ganglion cell when activated optogenetically, with respect to its normal photoreceptor response (fig.5.C). An analysis of the distribution of these indices showed that ON ganglion cells are consistently more sustained in their optogenetic responses (tested with the Wilcoxon rank-sum test, p-value below 1e-5; see methods), while the result is less clear for OFF cells.

We then checked if the optogenetic stimulation of AIIs preserved or disrupted the functional organization of retinal ganglion cells. Retinal ganglion cells can be classified in types: cells belonging to the same type are spatially arranged to cover uniformly the visual field, and produce similar responses to the same stimuli. To check if this organization was still present, we looked at correlations of the responses of pairs of retinal ganglion cells for both photoreceptor and optogenetic stimulations (fig. 5.D). If the functional arrangement is preserved in optogenetically reactivated retinas, these correlations should not vary significantly across these two conditions. We used the Pearson Correlation Coefficient to estimate the correlation of two cells responses. We looked at the correlations of all cell pairs with the same polarity, for both photoreceptor and optogenetics responses. We observed that for most of the ganglion cell pairs considered, these correlations were still present during optogenetic stimulation (fig. 5.D).

As a further estimation of the diversity of the responses, we calculated the dimensionality of the space of possible responses. For this we performed a PCA on the ensemble of all the ganglion cell average responses to the chirp stimulus. If all the cells responded the same way to the stimulus, the first principle component would explain all the variance in these responses. On the contrary, if all the responses are very different, it will take a lot of components to explain most of the variance. We found that, for both normal and reactivated retinas, we needed more than 6 components to explain more than 95% of the total variance in the response (fig. 5.E). This shows that our strategy is able to restore a large part of the diversity in the visual responses.

### AII stimulation in degenerated retinas restore ON and OFF responses

So far we worked with wild-type retinas where we could compare the same ganglion cells responding to photoreceptor stimulation and optogenetic stimulation of AII amacrine cells. However, in retinal dystrophies, the retinal network is rewired following the degeneration [Marc & Jones 2003]. Is AII stimulation still able to evoke responses after the rewiring imposed by degeneration? To test this, we performed the same experiments on rd1 mice. We obtained a similar expression pattern in rd1 mice as in wild type mice (fig. 6.A, B, C). We recorded the retinas at an age (3 months) when there are no measurable responses to light due to photoreceptor degeneration. We collected data accounting for a total population of 160 RGCs, measured their responses to light flashes, and consistently found both ON and OFF responses (fig. 6.D, E). This demonstrates that the rewiring of the network following degenerescence does not affect the ability of our strategy to restore diverse responses in ganglion cells.

**Figure 6:**
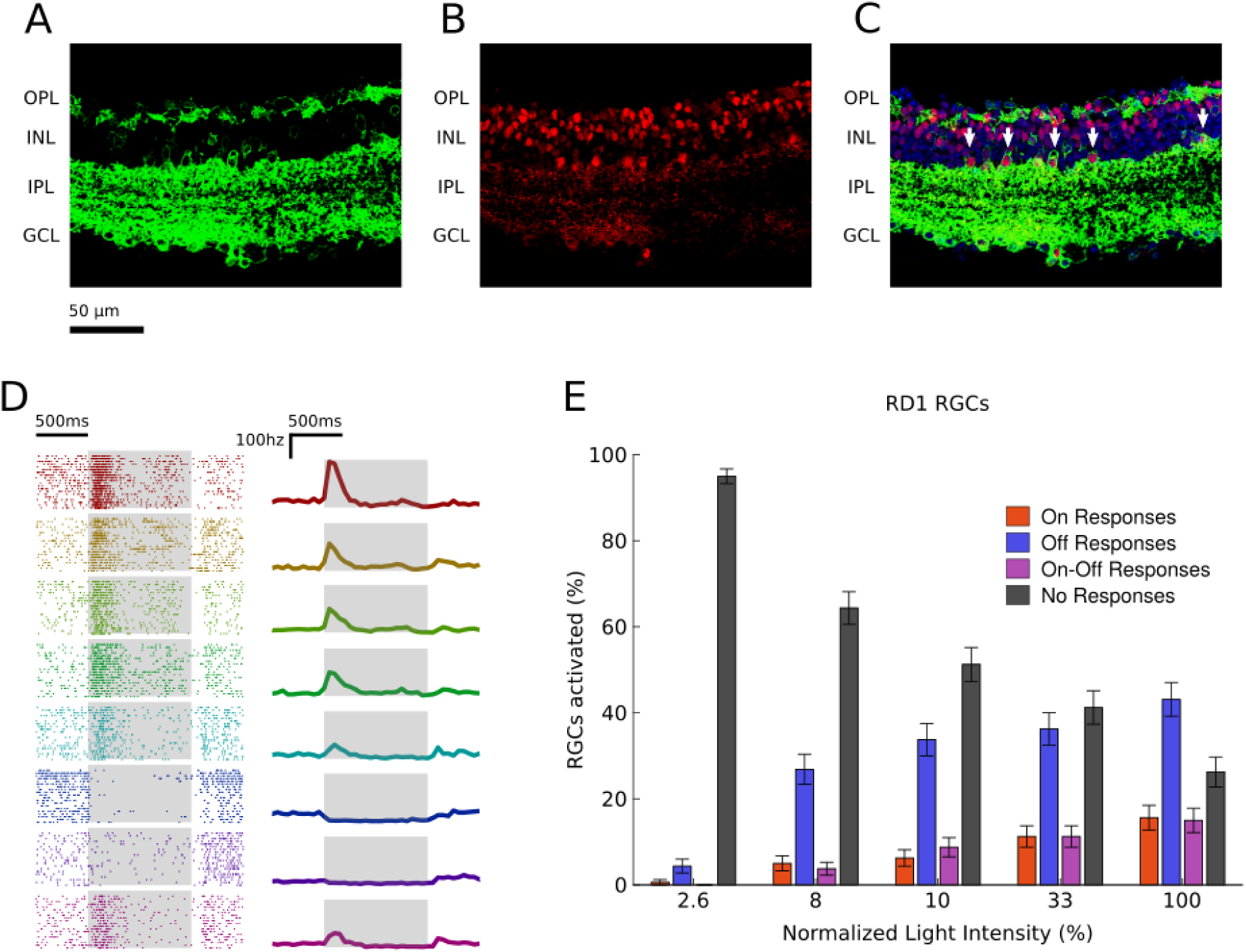
Optogenetic Activation of AIIs produces ON and OFF RGC responses also in dystrophic retinas. **A**,**B**,**C**. Cross section of a dystrophic mouse retina (rd1) showing ReachR expression (green, panels A and C) under HKamac promoter; co-localization with Prox1 antibody (red, panels B and C) indicates ReachR expression in AII amacrine cells. Expression under control of DAPI is shown in blue (panel C). AII amacrine cells are indicated with white arrows. Retinal layers are shown on the left: outer plexiform layer (OPL), inner nuclear layer (INL), inner plexiform layer (IPL), ganglion cell layer (GCL). **D**. Optogenetic responses of 10 representative retinal ganglion cells from dystrophic retinas to a series of flashes. Left: raster plots of the responses; each row (and color) represents a different cell; the stimulation period is highlighted in gray. Right: mean responses over trials. **E**. Activation of RGCs in dystrophic retina due to optogenetic stimulation. For each of five luminance levels we plot the percentage of retinal ganglion cells that responded to the optogenetic stimulation with pure ON (red), pure OFF (blue), ON-OFF (magenta) or no responses (black).

## Discussion

Here, we report a broadly applicable strategy that can be used to restore visual function in patients suffering from photoreceptor degeneration. We targeted, for the first time, AII amacrine cells thanks to a specific promoter and expressed high levels of microbial opsin to render them light sensitive. Our strategy has the advantage of being mutation-independent, and can potentially be used for different genotypes of retinal dystrophies. Compared to ganglion cell targeting, which is currently in clinical trials, our approach has the advantage of restoring a significant part of the retinal computations performed by a normal retina. In particular, we have shown that this strategy allows restoring both ON and OFF ganglion cell responses. We also observed diverse responses, presumably corresponding to the activation of several different cell types and pathways in the retinal circuit, similar to what happens in the normal retina.

Restoring this diversity of responses could be important for restoring visual perception. Previous studies on the primate retina [Schiller et al. 1986, Merigan et al. 1991] have shown that a selective impairment of retinal computations leads to specific deficits in visual perception. Inactivating ON cells in the macaque retina in vivo using pharmacology [Schiller et al. 1986] affected the ability of the macaque to detect light increase but did not affect its ability to detect light decrease. Recent work [Smeds et al. 2019] looking at responses of mouse RGCs and behavior at scotopic light levels, suggests that the mouse relies on the responses of ON RGCs to detect light increase, and on OFF RGCs to detect light decrease. A striking finding by [Smeds et al. 2019] was that mice engaged in a task where they had to detect light increase in darkness would not use the information available from OFF RGCs, even when they are more sensitive than ON RGCs. This strongly suggests that ON RGCs are used to detect light increase and OFF RGCs to detect light decrease, at least in scotopic conditions.

These previous results suggest that restoring retinal computations might be necessary for a blind patient to perform complex visual tasks. Previous studies have proposed alternative strategies targeting either “dormant” cones [Busskamp et al. 2010] or bipolar cells [Macé et al. 2015, Gaub et al. 2015, van Wyk et al. 2015, Cehajic-Kapetanovic et al. 2015]. In many retinal dystrophy patients, photoreceptors are not present in late stages, and bipolar cells can be partially degenerate [Francis et al. 2013]. Indeed, retinal ganglion cells and AII amacrine cells have been shown to be the most robust neuronal cell types during retinal degeneration. AII amacrine cells that are stable over a longer time [Strettoi et al. 2002], can therefore be used at the most advanced stages of degeneration.

However, ultimately, these advantages will have to be evaluated in primate models to attest translational feasibility. It remains to be seen if the same level of expression and specificity found here can be achieved in primate models. Nevertheless, our results show that targeting AII for optogenetic stimulation is a promising new avenue for vision restoration.

## Methods

### AAV Productions

Recombinant AAVs were produced by the plasmid cotransfection method, and the resulting lysates were purified via iodixanol gradient ultracentrifugation as previously described [Dalkara et al. 2013]. Briefly, 40% iodixanol fraction was concentrated and buffer exchanged using Amicon Centrifugal Filter Units (Millipore, Molsheim, France). Vector stocks were then titrated for DNase-resistant vector genomes by real-time qPCR relative to a standard.

### Animals And Intravitreal Injections

All experiments were done in accordance with Directive 2010/63/EU of the European Parliament. The protocol was approved by the Local Animal Ethics Committee of Paris 5 (CEEA 34). All mice used in this study were C3H/HeN (rd1 mice) or C57Bl6J mice (wild type) from Janvier Laboratories (Le Genest Saint Isle, France). For injections, mice were anesthetized with isoflurane (5% induction, 2% during the procedure). Pupils were dilated, and an ultrafine 30-gauge disposable needle was passed through the sclera, at the equator and next to the limbus, into the vitreous cavity. Injection of 1.5 μl stock containing 3.04 × 10e12 particles of AAV was made with direct observation of the needle in the center of the vitreous cavity.

### Immunohistochemistry

Mice were sacrificed in accordance with all animal facility protocols at the Institut de la Vision by CO2 inhalation and cervical dislocation. Eyes were removed and fixed 2-3 hrs in 4% formalin solution at RT. Eyes for sectioning were cryopreserved in 30% sucrose prior to embedding in Neg50 (Thermofisher, Waltham MA) and cut into 12 μm sections. Slides were warmed 10min, blocked 6-8hrs for GFP-tissue or 1hr for ReaChR-tissue in 6% NDS/1% BSA/0.5% triton/PBS at 4C, and incubated O/N in 50% block solution with anti-Prox1 1:500 (Biolegend, San Diego, CA; Rb), anti-ChAT 1:1K (Chemicon, Gt), washed 3×5 min PBS, incubated in secondary and DAPI 1:2K for 1-2hrs at RT, washed 3×5 min, and coverslipped in Permafluor (Thermofisher). The same procedure was used for flat-mounts but primary incubation was for 3 days and secondary incubation for 1 day using anti-GFP 1:500 (Abcam, Cambridge, UK; Chk).

### Multi-Electrode Array

MEA recordings were obtained from ex-vivo isolated flat mounted retinas of wild type mice and rd1 mice aged from 132 to 324 days. Mice were sacrificed by quick cervical dislocation, and eyeballs were removed and placed in Ames medium (Sigma-Aldrich, St Louis, MO; A1420) bubbled with 95% O2 and 5% CO2 at room temperature. Isolated retinas were placed on a cellulose membrane and gently pressed against a MEA (MEA256 iR-ITO; Multi-Channel Systems, Reutlingen, Germany), with the RGCs facing the electrodes. For wild type and rd1 recordings MEA with respectively 3 μm and 60 μm electrode spacing were used. Pharmacology was used to block photoreceptor to bipolar cell transmission with 5 μm L-AP4 and 1 μm ACET, followed by 200 μm CNQX and 10 μm CPP in some experiments to block all transmission to RGCs (Tocris, Bristol UK). All of the multi-electrode array recordings were processed with the software spyking-circus [Yger et al. 2018] to sort the recorded spikes and obtain templates of individual retinal ganglion cell responses.

### Light Stimulation

To study the responses of RGCs to optogenetic stimulation we used a flickering stimulus (referred to as flicker in the following) consisting of a series of white full-field flashes of one second duration, interleaved with one second intervals of darkness. Flashes were played both at low (∼0.1 μW cm^-2^) and high (∼2.8 μW cm^-2^) light intensities. To study the diversity of the cell responses, we used a chirp stimulus. This is a full field stimulus, lasting 25 seconds, designed to test the reaction of ganglion cells to changes in light intensity at different regimes of contrast and frequency. After a 1 s white flash, the light intensity varies at constant speed and increasing contrast for 10 s, and with constant contrast and increasing frequency for other 10 s.

### Detection of Ganglion Cell Responses

To check if a stimulus **s** evoked a response in a given ganglion cell, we did the following test. First, we considered a control window of 300 ms right before the stimulus onset. For a given cell **c**, we calculated the average spontaneous firing rate **r**_**ctrl**_^**c**,**s**^ and its standard deviation **σ**_**ctrl**_^**c**,**s**^ across repetitions in this time interval. We defined an activation threshold **T**^**c**,**s**^ as:

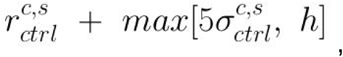

with **h** equal to 10hz. Then, we looked at a response window of 300 ms after the presentation of the stimulus (specifically, right after the stimulus onset for ON responses, and after the stimulus offset for OFF responses), and computed the peri-stimulus time histogram **psth**^**c**,**s**^ of these responses (time bin equal to 50 ms). We considered that there was a response if the mean response **psth**^**c**,**s**^ exceeded the activation threshold **T**^**c**,**s**^.

### Classification Of ON And OFF Ganglion Cells

We used the criterion above to classify ON and OFF retinal ganglion cells. We labeled as ON or OFF all the cells that responded respectively to the onset or to the offset of the flicker. Cells responding at both onset and offset (ON-OFF) and cells that were not responding at all were not labeled, and have not been considered in further analyses. For the excitatory opsin protocol (ReachR), we pooled data from three different experiments, obtaining a total population of 173 ON and 65 OFF ganglion cells. For the control protocol, we recorded from two different retinas, and identified a total of 113 ON and 94 OFF ganglion cells. For the inhibitory opsin protocol (gtACR), we collected data from a single experiment, and found 36 ON and 125 OFF ganglion cells.

### Analysis Of RGC Responses To Optogenetic Stimulation

To study the responses of RGC to the optogenetic stimulation, we displayed the flicker at low (∼0.1 μW cm^-2^) and high (∼2.8 μW cm^-2^) light intensities. Then, for each luminance level, we calculated the percentage of ON and OFF cells with detectable responses at both stimulus onset and offset. We did the same analysis to test the impact of both the excitatory and inhibitory opsins (respectively Reachr and gtACR1, fig. 2.D, 3.E, 4.C), and for all the control protocols with L-AP4, ACET and CNQX and CPP (fig. 2.E, 3.F). For the control experiments on rd1 mice, since the photoreceptor responses were not available, we could not preliminarily classify the ganglion cells as ON or OFF. As a consequence, we computed the percentages of RGCs activated on the entire population of cells, without any subdivision based on polarity (fig 6.E).

### Off-Target Expression Of Retinal Ganglion Cells

To identify the retinal ganglion cells directly expressing Reachr due to the expression leakage, we looked at the optogenetic responses to the flicker after application of CNQX and CPP. We labeled as affected by the leakage all those cells for which a detectable response (either at stimulus onset and/or offset) was detectable. We did not consider these RGCs for the sustained-transient analysis nor for the complexity analysis described below.

### Sustained-Transient Index

We defined a sustained-transient index **ST**^**c**,^ to assess which component is prevailing (transient or sustained) in the responses of a given ganglion cell **c** to a certain stimulus **s**. We considered two response windows subsequent to the presentation of the stimulus: a first one capturing transient responses (from 0 ms to 300 ms after stimulus onset for ON cells, and after stimulus offset for OFF cells) and a second one for sustained responses (from 300 ms to 600 ms after onset for ON cells and after offset for OFF cells). We computed the peri-stimulus time histogram of the responses (time bin equal to 50 ms) on both windows: **psth**_**trans**_^**c**,**s**^ representing the transient component, and **psth**_**sust**_^**c**,**s**^ representing the sustained component. We then computed the sustained-transient index **ST**^**c**,**s**^ as the ratio:

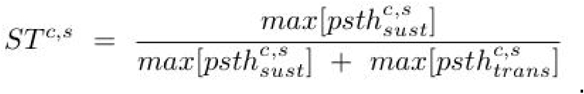

An index **ST**^**c**,**s**^ close to 1 entails that the cell **c** produces a sustained response to the stimulus **s**. Conversely, an index close to 0 means the response is predominantly transient.

### Sustained-Transient Analysis

We used the **ST** index defined above to compare the transience of RGC responses to photoreceptors and optogenetic stimulations respectively. For this analysis, we only considered cells with detectable responses to both the photoreceptor and optogenetic responses. We also excluded all the cells for which optogenetic responses showed a different polarity with respect to the photoreceptor responses, and all those cells affected by the expression leakage, leaving us with a total of 48 good cells. We then calculated the sustained-transient index described above for all cells in both optogenetic and photoreceptor responses, and compared the distribution of the index under the two conditions using the Wilcoxon rank sum test. We found that, for the population of ON ganglion cells, these two distributions are significantly different, with the responses to optogenetic stimulation being more sustained (the test rejects the null hypothesis that the two distributions have equal medians with p-val < 1e-5). For the population of OFF cells, we did not observe a significant difference among the two distributions (fig 5.B).

To consolidate this result at single cell level, we computed for each cell a comparative index **ΔST**, defined as the difference of the ST indices computed respectively on the photoreceptor and optogenetic responses:

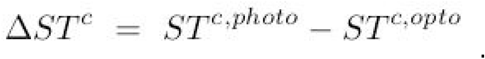

This index has value close to 0 if the photoreceptor and optogenetic responses are similar in transience, value close to 1 if the cell has an optogenetic response more transient with respect to its normal photoreceptor responses, and value close to -1 if its optogenetic responses are more sustained. We looked at the distribution of this index for both ON and OFF ganglion cells (fig 5.C), and observed that the vast majority of ON retinal ganglion cells (31 cells out of 34) has a negative relative index, indicating that ON ganglion cells tend to lose their transient component when activated optogenetically through the AII pathway.

### Analysis Of Complexity Of The Responses

To assess the complexity of RGC responses we relied on principal component analysis (PCA). We computed the principal components of the ganglion cell responses to the chirp stimulus for both photoreceptor and optogenetic stimulation, and compared the number of components needed to explain different percentages of variance. We applied the selection criteria used for the sustained-transient analysis described above: we only kept cells consistently responding to both photoreceptor and optogenetic stimulation, and excluded all the cells featuring off-target expression, for a total of 40 good cells. As we did not want to account for inter-experimental variability, we ran the analysis independently for each of the three experiments. To make the results comparable, we needed to keep the population size constant across the different experiments. As a consequence, we ran the principal component analysis on each experiment on a sampled subpopulation of fixed size (30 cells). We repeated this procedure 100 times, and for each experiment we computed the average curve showing the variance explained by each number of principal components under both normal and optogenetic conditions (fig. 5.E). We then obtained the final results by averaging the curves across all three experiments (opaque lines in figure fig. 5.E).

## Supporting information

Supplemental Material 1: hkamac-reachr sequence

## Acknowledgements

We thank the Paris Vision Institute core facilities (Animal facility platform, Histology platform, Imaging platform) and particularly Camille Robert of the vector core facility for producing the AAVs. This work was supported ERC Starting Grant (REGNETHER 639888/DD), the Centre National de la Recherche Scientifique (CNRS), the Institut National de la Santé et de la Recherche Médicale (INSERM), Labex-Lifesenses (D.D., J.D.), Sorbonne Université, INSERM, LabEx LIFESENSES (ANR-10-LABX-65), IHU FOReSIGHT (ANR-18-IAHU-01), Paris Ile-de-France Region under « DIM Thérapie génique » initiative. Foundation Fighting Blindness (Program Project Award).

## Author Contributions

Designed experiments: H.K., E.O., F.T., D.D., O.M.

Performed experiments: H.K., E.O., M.R., M.D.

Analyzed the data: F.T., P.Y., O.M.

Wrote the paper: H.K., E.O., F.T., D.D., O.M.

